# Spatial transcriptomics reveals site-specific cellular and metabolic heterogeneity in bladder carcinoma in situ

**DOI:** 10.64898/2026.08.27.741603

**Authors:** Tyler D. Myers, Amirali Salmasi, Margaret F. Meagher, Sarah Azari, Sonya Donato, Iveta Kalcheva, Se Jin Song, Haiyan Zhang, Kit Yuen, Aditya Bagrodia, Tyler F. Stewart, Michael Liss, Andrew Bartko

**Affiliations:** Center for Microbiome Innovation, Jacobs School of Engineering, University of California San Diego, La Jolla, CA, USA; Department of Urology, University of California San Diego, La Jolla, CA, USA; Department of Pathology, University of California San Diego, La Jolla, CA, USA; Department of Medicine, Division of Hematology-Oncology, University of California San Diego, CA, USA; Department of Pediatrics, University of California San Diego, La Jolla, CA, USA; Shu Chien-Gene Lay Department of Bioengineering, University of California San Diego, La Jolla, CA, USA

**Keywords:** Bladder cancer, Carcinoma in situ, Spatial transcriptomics, Intratumoral heterogeneity, Tumor microenvironment, Flux balance analysis, BCG-unresponsive, Non-muscle-invasive bladder cancer

## Abstract

Bladder carcinoma in situ (CIS) is a multifocal, non–muscle-invasive disease with a high risk of progression to muscle-invasive cancer. Current management strategies are often guided by genomic profiling of single tumor samples, which incompletely capture tumor heterogeneity and may contribute to treatment failure. In particular, the multifocal nature of CIS raises uncertainty regarding the uniformity of genomic, immunologic, and microenvironmental features across anatomically distinct sites within the same patient. To address this, we performed spatial transcriptomic profiling of CIS-containing tissue from four anatomically distinct sites within a single individual. Unsupervised clustering with marker-based annotation, integrated with metabolic inference, identified epithelial tumor populations alongside stromal, immune, and smooth muscle compartments. While key cellular states were conserved, their spatial organization and relative abundance varied by site. Metabolic analysis further revealed region-specific microenvironments shaped by local cellular architecture. These findings indicate that both cellular composition and metabolic activity are spatially structured. Collectively, these results demonstrate that CIS exhibits significant intra-patient heterogeneity not captured by single-site profiling. These findings require validation in larger cohorts but support multi-region sampling could help improve risk stratification, biomarker development, and prediction of response to intravesical therapies, with potential implications for more personalized treatment strategies.

## Main

Bladder carcinoma in situ (CIS) is a multifocal, high-grade malignancy associated with substantial risk of progression to muscle-invasive disease in untreated patients or in those who fail intravesical therapy^1^. Management of CIS remains challenging because of its multifocal nature and variability in pathologic reporting, with some pathologists describing it as a distinct entity and others as a concurrent “shoulder lesion” adjacent to high-grade papillary tumors. CIS also serves as a key eligibility criterion in clinical trials targeting both BCG-naïve and BCG-unresponsive non–muscle-invasive bladder cancer (NMIBC), yet its underlying biology remains incompletely understood^2^. Current risk stratification and biomarker development rely heavily on genomic and immunologic profiling derived from single tumor samples, which implicitly assume tumor homogeneity and may fail to capture clinically relevant intratumoral heterogeneity^3,4^.

Here, we applied multi-region spatial transcriptomics within a single patient to test the hypothesis that anatomically distinct bladder CIS regions harbor divergent cellular, spatial, and metabolic programs that are not captured by single-site sampling. Across four anatomically distinct bladder regions (the bladder dome, bladder trigone, prostatic urethra, and distal ureter) in a patient who underwent radical cystectomy for BCG-unresponsive NMIBC, we profiled 109,600 spatial elements at 50 µm resolution (derived from 500 nm capture features) spanning up to 1 cm^2^ of tissue. This revealed conserved cellular programs alongside marked site-specific differences in composition and spatial architecture (**Figure 1a–c**).

**Figure 1.**
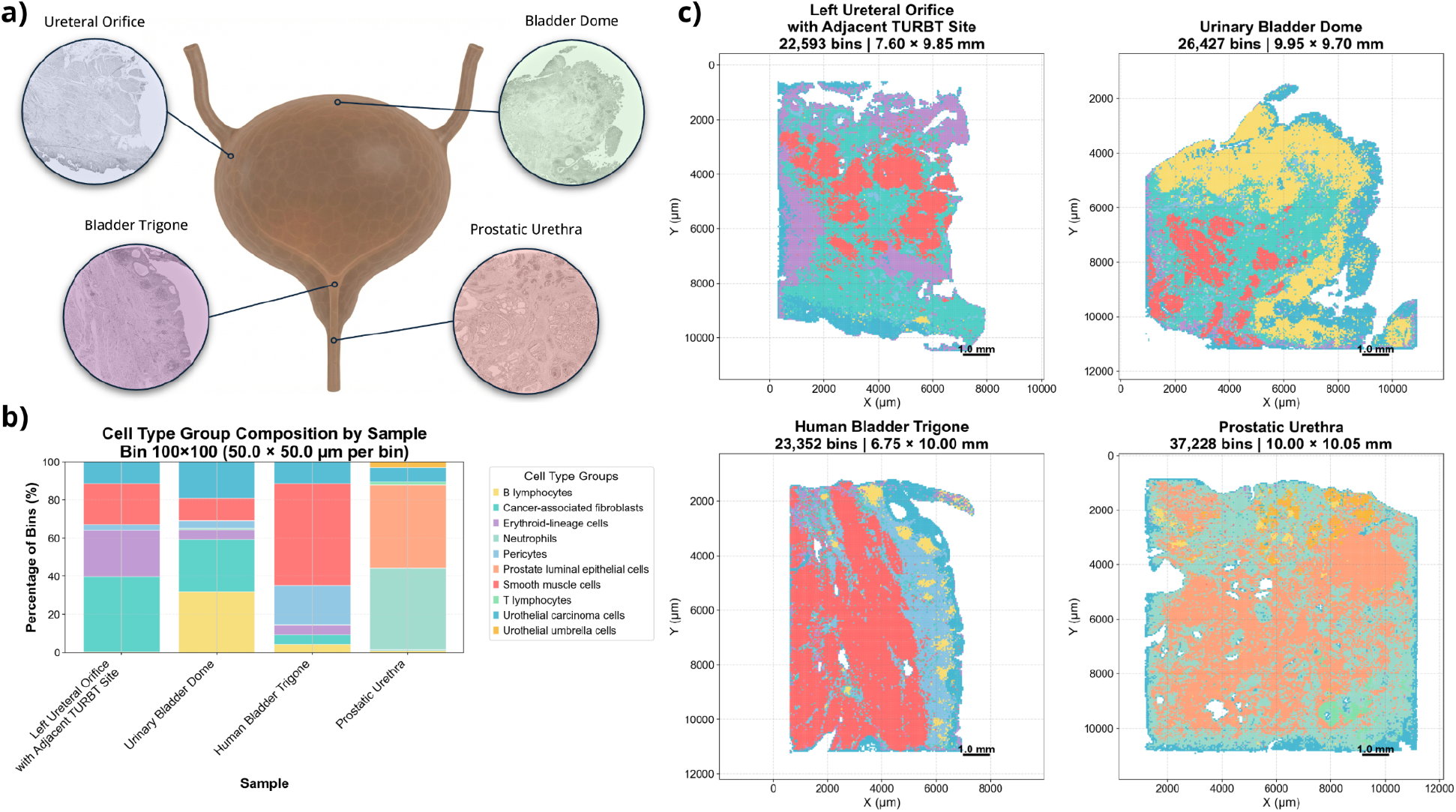
Spatial transcriptomics profiles and cellular composition of urothelial carcinoma show intratumor heterogeneity. **a)** Overview of the four sites sampled for spatial transcriptomics analysis. **b)** Per-site tissue cell type composition using the inferred cell type labels derived from clustering and up-regulated gene analysis. **c)** Spatial transcriptomics profiles for each of the four bladder sites, colored by the cell type label.

Unsupervised clustering with marker-based annotation identified distinct urothelial carcinoma–associated epithelial populations, including LCN2^+^/KRT13^+^ and TSPAN12-high subtypes, alongside normal urothelial (PIGR^+^ umbrella cells) and region-specific anatomical elements such as KLK3^+^/MSMB^+^ prostatic luminal epithelium (**Supplementary Table S1**). These epithelial compartments coexisted with smooth muscle cells (CARMN, MYH11), pericytes (ENPEP, NTRK3), cancer-associated fibroblasts (CAFs; SERPINE1, GREM1, MFAP5), and infiltrating immune populations (MS4A1^+^ B cells, RNF128^+^ T cells, LTF^+^ neutrophils, and HBA2^+^ erythroid-lineage cells; **Supplementary Figure S2**). Cellular composition varied substantially by anatomical site. The bladder trigone and ureteral sites were enriched for fibroblast- and smooth muscle–associated programs (22% and 38% of cellular composition, respectively), consistent with mixed tumor–stroma interfaces. In contrast, the prostatic urethra was dominated by prostate luminal epithelial cells (43%), while the bladder dome exhibited a higher immune cell representation (32%), indicating regionally distinct immune niches (**Figure 1b–c**). These findings highlight substantial intra-patient heterogeneity that would not be captured by single-site sampling. Beyond regional cell-type composition, cellular compartments exhibited spatial clustering rather than uniform mixing (**Figure 1c**). Urothelial carcinoma cells formed discrete epithelial nests bordered by fibrovascular stroma and localized immune infiltrates, creating compartmentalized functional niches^5^. The degree of tumor-stroma intermixing likely acts as a physical barrier that promotes local immunosuppression, restricts T-cell infiltration, and may influence responses to immunotherapy.

To extend beyond marker-based annotation, we applied spatial flux balance analysis^6^ to infer per-bin metabolic activity. Flux maps for the ureteral site are shown in detail (**Figure 2b**), with expanded maps for all four anatomical sites provided in **Supplementary Figure S1**. Flux maps demonstrated spatially heterogeneous metabolism, including region-specific glucose consumption, lactate production, and variable pyruvate import into mitochondria, consistent with locally organized oxidative phosphorylation rather than a uniform Warburg phenotype^7^.

**Figure 2.**
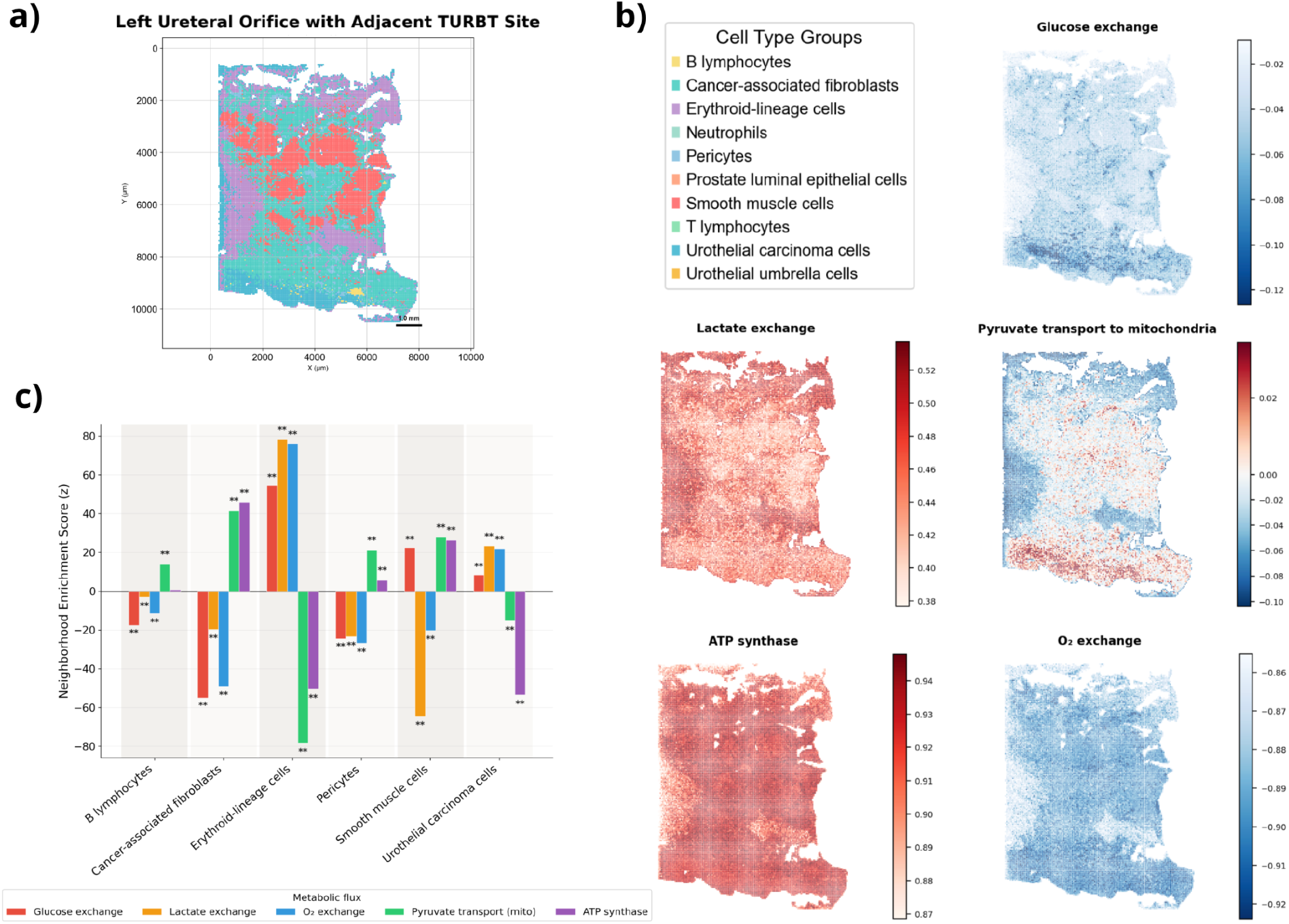
Flux balance analysis and neighborhood enrichment co-occurrence analysis highlight intratissue heterogeneity. **a)** Left Ureteral Orifice with Adjacent TURBT Site spatial transcriptomics profiles colored by cell type. **b)** Metabolic flux for Glucose, Lactate, Pyruvate transport to mitochondria, ATP synthase, and O_2_ exchange. In the case of exchange reactions, negative values correspond to consumption of the metabolite, positive values to production. **c)** Neighborhood enrichment score for the cell types in the Left Ureteral Orifice with Adjacent TURBT Site sample, with bar colors indicating the metabolic flux.

Permutation-based spatial enrichment within the ureteral site linked specific cell populations to distinct metabolic environments. Cancer-associated fibroblasts (CAFs) were significantly enriched in regions of high glucose and oxygen consumption (Glucose: Z=−54.93, p<0.001; Oxygen: Z=−48.98, p<0.001; **Figure 2c**), consistent with a metabolically competitive niche expected to restrict glycolysis-dependent effector T-cell activity and to create localized immunosuppression^8^. These spatially restricted immunosuppressive niches have important clinical implications, as single-site biopsies may fail to capture regional heterogeneity and risk biomarker misclassification for immunotherapies and antibody-drug conjugates. Erythroid-associated compartments showed enrichment for lactate exchange (Z=78.33, p<0.001) and reduced pyruvate import into mitochondria (Z=−78.15, p<0.001), consistent with the obligate anaerobic glycolysis of erythrocytes and serving as an internal positive control for the inference pipeline. Although demonstrated in a representative region, this framework provides a scalable approach to link spatial cellular architecture with inferred metabolic programs across anatomical sites.

Collectively, these findings demonstrate that anatomically distinct CIS regions within a single patient harbor substantial heterogeneity in cellular composition, spatial organization, and metabolic activity. Given that current risk stratification and therapeutic decision-making rely heavily on single-tumor profiling, these results highlight a critical gap in our understanding of CIS biology^9,10^. Given these observations are hypothesis-generating and derived from a single case, they require validation in larger, multi-patient cohorts and support the need for multi-region sampling strategies to more accurately define clinically relevant heterogeneity and improve biomarker development and treatment selection.

## Methods

### Sample Preparation and Sequencing

FFPE blocks of tissue was obtained from four distinct anatomical bladder sites (left ureteral orifice with adjacent TURBT site, bladder trigone, urinary bladder dome, and prostatic urethra) from one patient. Sections of the blocks were processed using the STOmics Stereo-seq workflow following manufacturer protocols for sectioning and chip mounting (STUM-SP003), transcriptomics preparation (STUM-TT004), library preparation (STUM-LP001), and sequencing CSS-00037). Briefly, a section of the FFPE block was cut to a thickness of 5μm and adhered to the Stereo-seq chip, which then underwent deparaffinization, imaging, decrosslinking, fixation, permeabilization, reverse transcription, cDNA release and collection, and finally cDNA purification and amplification. Libraries were prepared from the amplified cDNA and sequenced on a DNBSEQ-T7 instrument.

### Spatial Data Processing and Cell Clustering

Primary alignment and count matrix generation were executed via the SAW v8.1.3 pipeline^11^ using default settings and T2T-CHM13 as the reference genome for alignment and annotation. Spatial coordinates were binned at a 100 × 100 capture unit resolution (50 µm/bin). Bins were quality-filtered by requiring a minimum of 50 counts per bin and including genes detected in at least 10 bins. Filtered gene expression matrices were normalized to a total count of 10,000 per bin and log1p-transformed. The genes most likely to discriminate between cell types were considered those with the highest variance-to-mean ratio. Principal component analysis (PCA) was used on the top 20,000 most variable genes with mean expression >0.1 to reduce dimensionality to 50 components, followed by nearest-neighbor graph construction in the PCA embedding space. Unsupervised community detection was performed using the Leiden algorithm at a fixed resolution of 1.403, resolving 13 clusters across the four combined samples. Cluster marker genes were identified using Wilcoxon rank-sum differential expression testing, and cell types were annotated using a curated marker dictionary informed by the publications listed in Table S1.

### Spatial Metabolic Flux Analysis

To model localized metabolic activity, spatial flux variability and sampling were conducted using the Spatial Flux Balance Analysis (spFBA) framework^12^. Gene expression data were mapped to the human genome-scale metabolic reconstruction model *ENGRO2*^13^ using COBRApy. As recommended in Galuzzi et al., 2022^14^, gene expression measurements across spatial coordinates were denoised using Markov Affinity-based Graph Imputation of Cells (MAGIC)^15^ prior to computing Reaction Activity Scores (RAS) for each spatial bin. Spatial distribution of fluxes for 12 candidate reactions—including mitochondrial transport of glucose, lactate, pyruvate, ATP synthase activity, and oxygen exchange—were evaluated across tissues.

### Cell-Type and Metabolic Flux Neighborhood Enrichment

To evaluate the spatial relationship between cell-type localization and metabolic microenvironments, spatial neighborhood enrichment testing was performed. A k-nearest-neighbor (KNN) spatial graph was constructed directly from bin spatial coordinates. Permutation testing was applied to determine whether specific cell types exhibited significantly elevated or depleted flux levels within the same bin (within-element enrichment) or across immediate spatial neighbors relative to random expectation. Multiple testing correction was applied using the Benjamini–Hochberg False Discovery Rate (FDR) procedure. Group-level differences in flux distributions across cell types were validated using Kruskal–Wallis omnibus tests followed by post-hoc Dunn’s tests with FDR adjustment.

## Acknowledgments

This work was supported by grants from the UC San Diego Academic Senate awarded to A.S. We are also deeply grateful to the patient who generously consented to provide tissue samples for this study, making this work possible.

## Declaration of Generative AI and AI-assisted technologies in the writing process

During the preparation of this work, the authors used OpenAI Deep Research to help find and summarize the genes associated with cell types from the existing literature (see Supplementary Table S1). The authors reviewed and edited the output as needed and take full responsibility for the content of the published article.

