## Supplemental Figures and Table for "Spatial transcriptomics reveals site-specific cellular and metabolic heterogeneity in bladder carcinoma in situ"

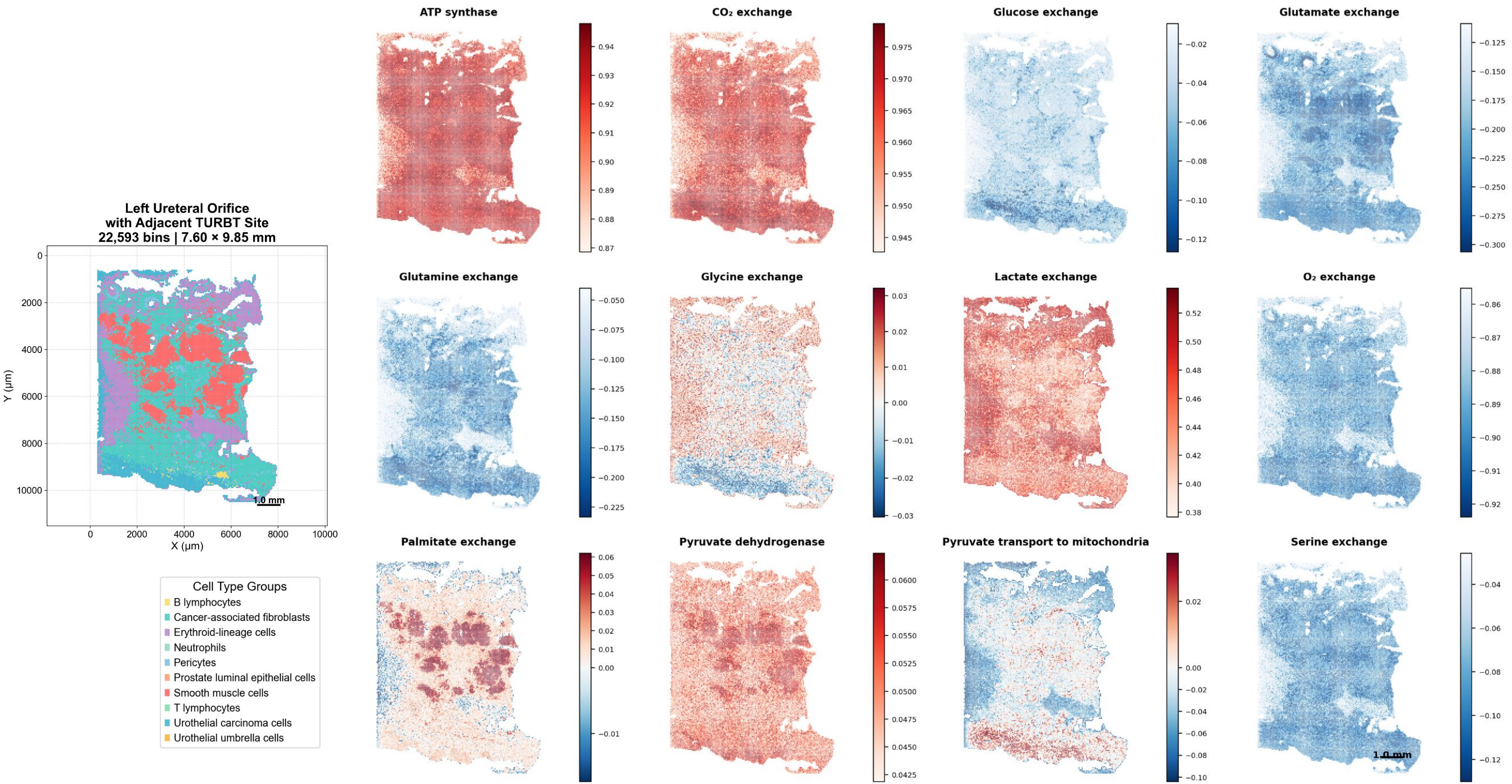


**Figure S1a.** Flux balance analysis of the spatial transcriptomics reveals cell type specific metabolisms throughout various bladder tissues: a) Left Ureteral Orifice with Adjacent TURBT Site, b) Urinary Bladder Dome, c) Bladder Trigone, and d) Prostatic Urethra. **Left)** Spatial transcriptomics profiles colored by inferred cell type. **Right)** Metabolic flux for various activity types. In the case of exchange reactions, negative values correspond to consumption of the metabolite, positive values to production.

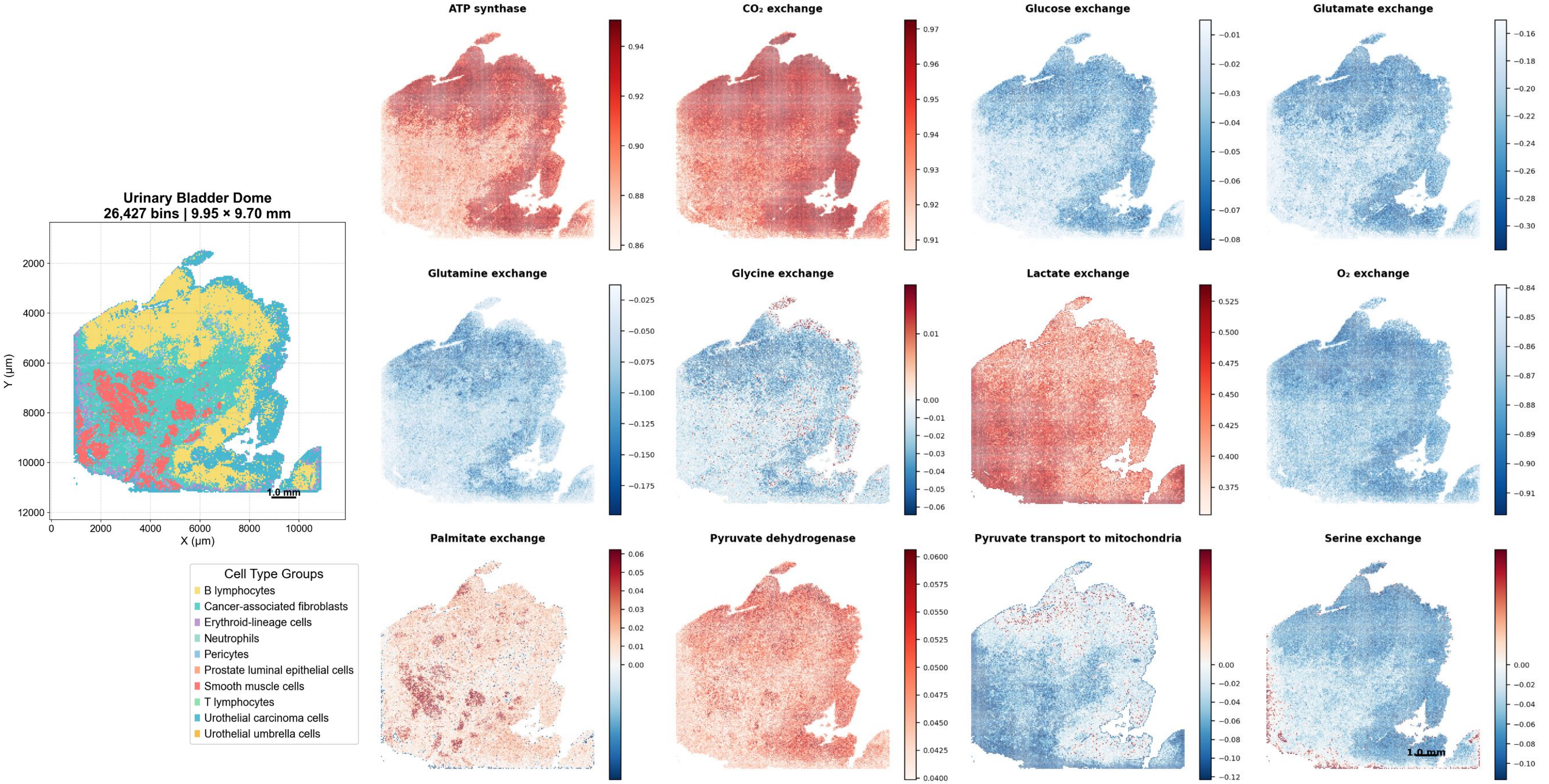

Urinary Bladder Dome
26,427 bins | 9.95 × 9.70 mm
Y (µm)
X (µm)
1.0 mm
Cell Type Groups
B lymphocytes
Cancer-associated fibroblasts
Erythroid-lineage cells
Neutrophils
Pericytes
Prostate luminal epithelial cells
Smooth muscle cells
T lymphocytes
Urothelial carcinoma cells
Urothelial umbrella cells
ATP synthase
CO₂ exchange
Glucose exchange
Glutamate exchange
Glutamine exchange
Glycine exchange
Lactate exchange
O₂ exchange
Palmitate exchange
Pyruvate dehydrogenase
Pyruvate transport to mitochondria
Serine exchange


**Figure S1b.**

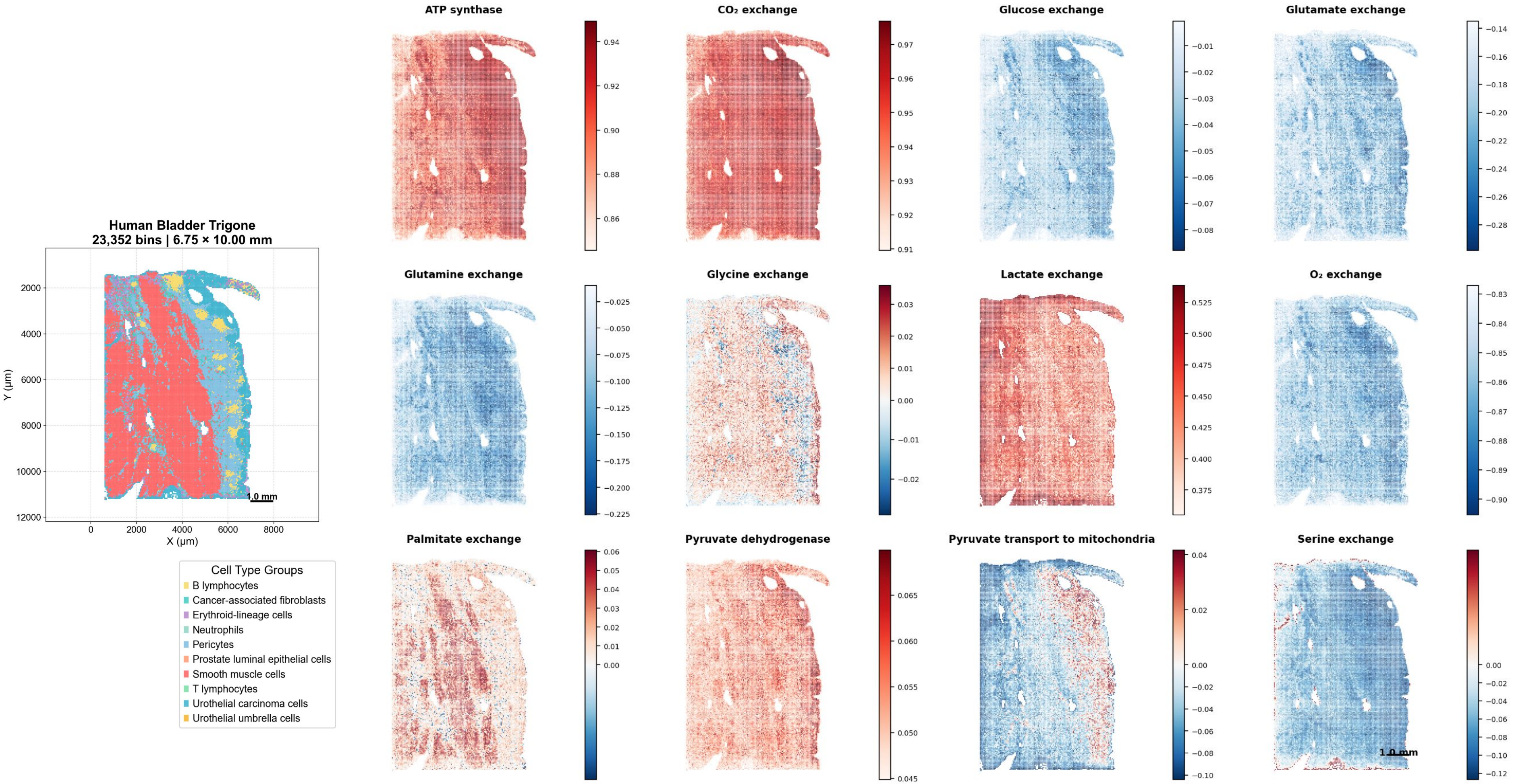
Human Bladder Trigone
23,352 bins | 6.75 × 10.00 mm
Y (μm)
X (μm)
1.0 mm
Cell Type Groups
B lymphocytes
Cancer-associated fibroblasts
Erythroid-lineage cells
Neutrophils
Pericytes
Prostate luminal epithelial cells
Smooth muscle cells
T lymphocytes
Urothelial carcinoma cells
Urothelial umbrella cells
ATP synthase
CO₂ exchange
Glucose exchange
Glutamate exchange
Glutamine exchange
Glycine exchange
Lactate exchange
O₂ exchange
Palmitate exchange
Pyruvate dehydrogenase
Pyruvate transport to mitochondria
Serine exchange

**Figure S1c.**

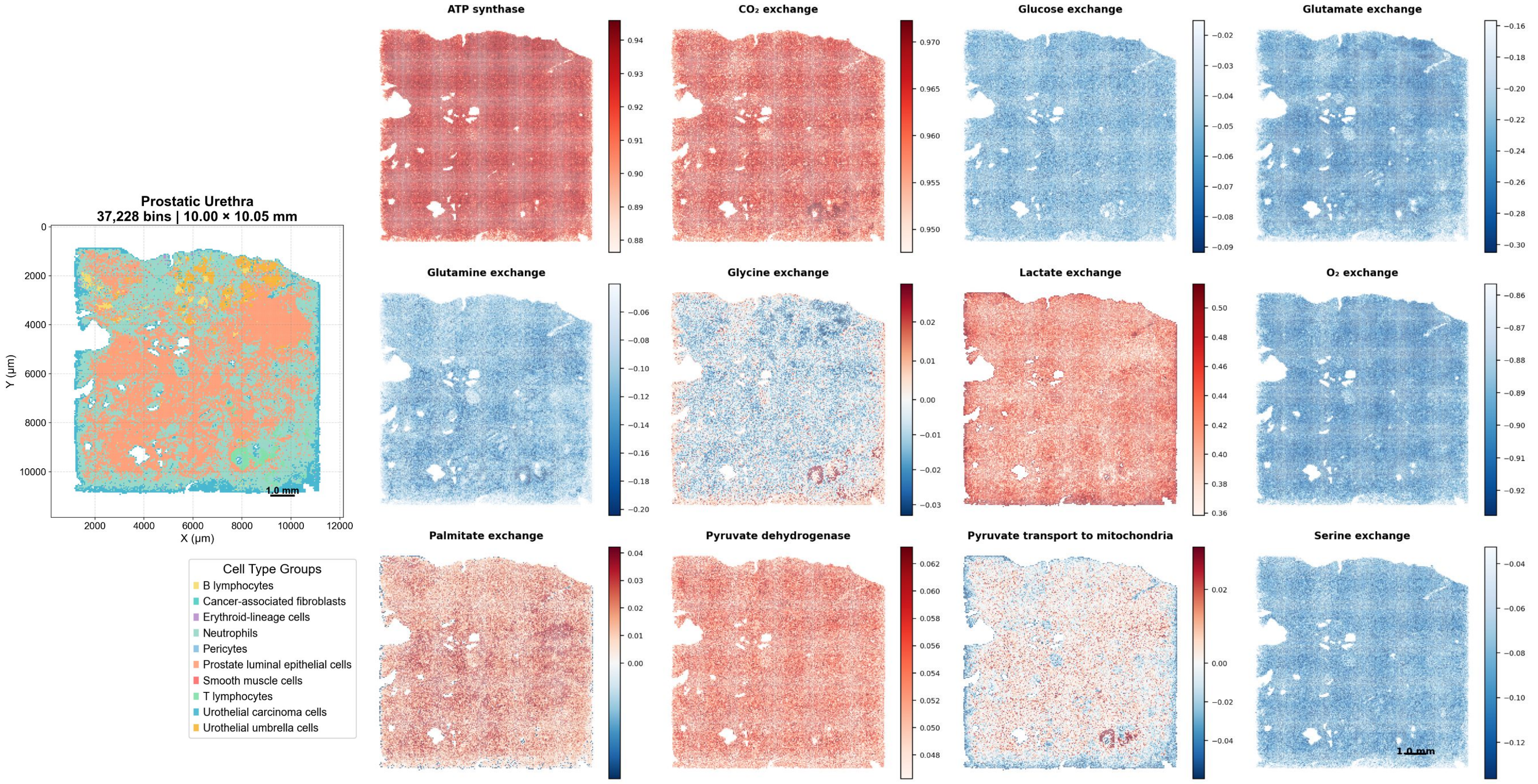

Prostatic Urethra
37,228 bins | 10.00 × 10.05 mm
Y (µm)
X (µm)
1.0 mm
Cell Type Groups
B lymphocytes
Cancer-associated fibroblasts
Erythroid-lineage cells
Neutrophils
Pericytes
Prostate luminal epithelial cells
Smooth muscle cells
T lymphocytes
Urothelial carcinoma cells
Urothelial umbrella cells
ATP synthase
CO₂ exchange
Glucose exchange
Glutamate exchange
Glutamine exchange
Glycine exchange
Lactate exchange
O₂ exchange
Palmitate exchange
Pyruvate dehydrogenase
Pyruvate transport to mitochondria
Serine exchange
1.0 mm


**Figure S1d.**

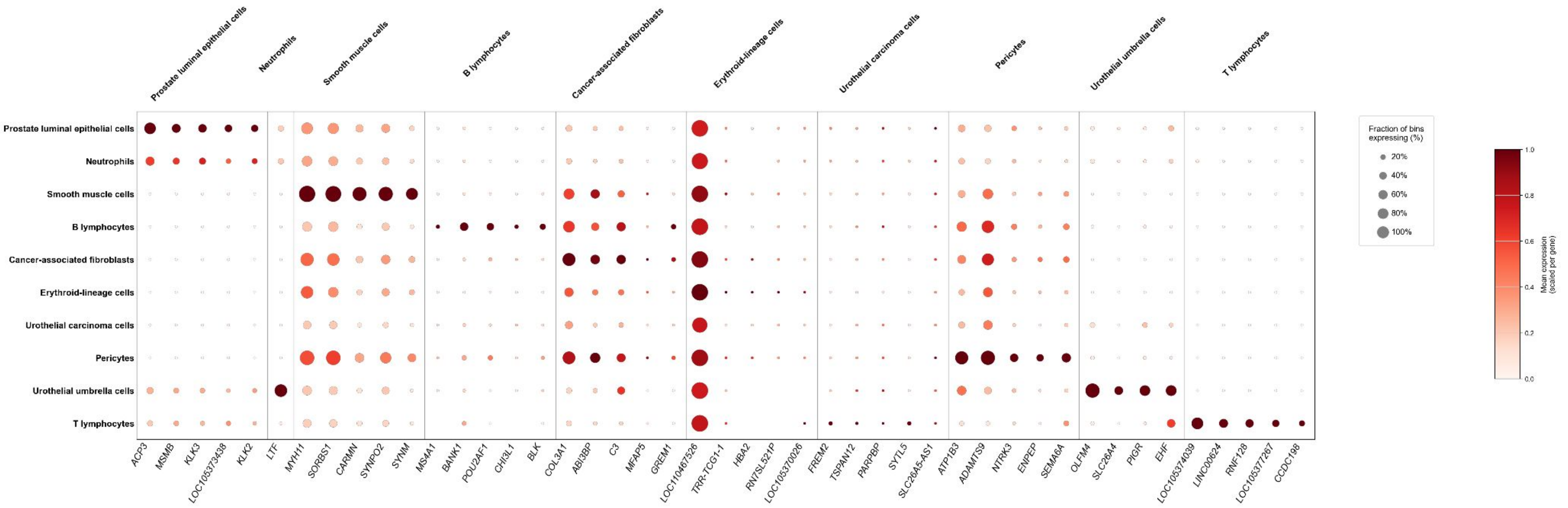


**Figure S2.** Expression of genes across inferred cell types. Dot size shows the percentage of the bins expressing the gene and color shows the relative expression level.

| Cluster | Upregulated genes | Inferred cell type | Supporting PubMed IDs |
|---|---|---|---|
| Cluster 0 | ACP3, MSMB, KLK3, LOC105373438, KLK2 | Prostate luminal epithelial cells (prostatic gland contamination/metastatic component) – indicated by prostate-specific secretory genes KLK2/KLK3 (PSA) and MSMB | 35763641 |
| Cluster 1 | LTF | Neutrophils – Lactotransferrin (LTF) is a prominent component of neutrophil secondary granules | 16261255 |
| Cluster 2 | CARMN, SORBS1, ACTG2, RBFOX3, TPM1 | Smooth muscle cells (bladder wall muscle/perivascular SMC) – express contractile proteins (e.g. ACTG2, TPM1) and the SMC-specific lncRNA CARMN | 34694145 |
| Cluster 3 | MS4A1, BANK1, POU2AF1, CHI3L1, BLK | B lymphocytes (tumor-infiltrating B-cells) – MS4A1 encodes the B-cell marker CD20, and POU2AF1 (BOB1/OCA-B) is a B-cell-specific transcription coactivator | 39684762, 40786043 |
| Cluster 4 | EMP1, LOC110467527, NFATC2, ADAMTS9, SERPINE1 | Cancer-associated fibroblasts (activated stromal fibroblasts) – characterized by TGFβ-induced and matrix-remodeling genes (e.g. SERPINE1/PAI-1) often upregulated in desmoplastic tumor stroma | 35267539 |
| Cluster 5 | MYH11, SYNPO2, SYNM, COL3A1, ABI3BP | Smooth muscle cells (detrusor muscle/myofibroblastic stroma) – marked by smooth muscle contractile genes like MYH11 (muscle myosin heavy chain) and collagen (COL3A1) production in the stroma | 34694145, 38890455 |
| Cluster 6 | C3, MFAP5, GREM1, LOC110467526, TRR-TCG1-1 | Cancer-associated fibroblasts (secretory/inflammatory CAF subtype) – defined by ECM and signaling factors (MFAP5, GREM1) produced by tumor fibroblasts | 31190277, 40940956 |
| Cluster 7 | HBA2, RN7SL521P, LOC105370026 | Erythroid-lineage cells (red blood cell cluster, e.g. reticulocytes) – high expression of hemoglobin α (HBA2) is specific to erythrocytes/precursors | 35514346 |
| Cluster 8 | LCN2, KRT13, DUOX2, TMPRSS4, MIR31HG | Urothelial carcinoma cells (squamous-/basal-like tumor cells) – exemplified by LCN2 overexpression, which correlates with high-grade, invasive bladder cancers (KRT13^+ differentiated tumor subset) | 41363723, 36989746 |
| Cluster 9 | ATP1B3, NTRK3, ENPEP, SEMA6A, FREM2 | Pericytes (vascular smooth muscle progenitors) – marked by ENPEP (aminopeptidase A) and NTRK3, which are enriched in vascular SMC/pericyte populations | 39661628, |
| Cluster 10 | TSPAN12, PARPBP, SYTL5, SLC26A5-AS1 | Urothelial carcinoma cells (minor epithelial subpopulation) – TSPAN12^high tumor epithelial cells, potentially engaged in stromal interaction (TSPAN12 is implicated in cancer–fibroblast crosstalk and Wnt signaling in tumors) | 25512506 |
| Cluster 11 | OLFM4, SLC26A4, PIGR, EHF | Urothelial umbrella cells (differentiated superficial urothelium) – characterized by PIGR expression on superficial epithelial cells for IgA transcytosis and differentiation factors like EHF | 1759330, 38890455 |
| Cluster 12 | LOC105374039, LINC00624, RNF128, LOC105377267, CCDC198 | T lymphocytes (tumor-infiltrating T-cells) – RNF128 (GRAIL) is a T-cell–specific E3 ubiquitin ligase involved in limiting T-cell activation and cytokine production | 20493730 |

**Table S1.** Upregulated genes for each cluster. From a broad literature search, cell types are inferred based on top upregulated genes, with supporting citations included. Literature sourced and summarized using OpenAI Deep Research without prior information pertaining to the location of the tissue sample. Citations and literature relevance were human verified and curated. Note: Cluster 10 may be an edge effect and all cell type labels should be reevaluated with a positive control or reference expression database for improved labeling.
